# Drivers and implications of the extreme 2022 wildfire season in Southwest Europe

**DOI:** 10.1101/2022.09.29.510113

**Authors:** Marcos Rodrigues, Àngel Cunill Camprubí, Rodrigo Balaguer-Romano, Julien Ruffault, Paulo M Fernandes, Víctor Resco de Dios

## Abstract

Wildfire is a common phenomenon in Mediterranean countries but the 2022 fire season has been extreme in southwest Europe (Portugal, Spain and France). Burned area has exceeded the 2001-2021 median by a factor of 52 in some regions and large wildfires started to occur in June-July, earlier than the traditional fire season. These anomalies were associated with record-breaking values of fuel dryness, atmospheric water demand and pyrometeorological conditions. For instance, live fuel moisture content was below the historical minima for almost 50% of the season in some regions. Wildfire impacts are primarily social and economical in these fire-prone landscapes, but they may prompt large-scale degradation if this anomaly becomes more common under climate change, as is expected. As climate changes intensify, we can expect this to become the new normal in large parts of the continent. Climate change is already here and delaying fuel management will only worsen the wildfire problem. Here we provide a preliminary though comprehensive analysis of 2022’s wildfire season in southwest Europe (Portugal, France and Spain).

## 1. Introduction

Wildfires are a natural phenomenon in Mediterranean countries, playing a key role in the conservation of landscapes and the dynamics of forest communities (Pausas et al., 2017). When the natural regimes of fire are altered due to increases in fire intensity and severity from global change, fires can threaten both the environment and society (Cochrane & Bowman, 2021; Moreira et al., 2020; Wunder et al., 2021). The confluence of fire exclusion and fuel build-up over the last decades, together climate change have set the conditions for unprecedented situations, already witnessing earlier and longer wildfire seasons (AghaKouchak et al., 2020; Moreira et al., 2020).

The 2022 fire season in southwest Europe (Portugal, Spain and France) has drawn considerable international attention due to the large extent of burned area, as 469,464 ha have burned at the time of writing (28^th^ September 2022), which is nearly 3 times higher than the 2006-2021 annual mean (173,415 ha; EFFIS dataset, explained below in methods). The season coincided with the chained irruption of several heat waves, which have appeared earlier than usual breaking temperature records in several countries like Spain or France (C3S, 2022), leading to record-breaking wildfire events. This season could thus potentially act as a spyglass into what the “new normal” will look like under climate change in forthcoming years. Understanding the processes and underlying drivers of these unprecedent events are crucial to mitigate and building fire-adapted and resilient landscapes and communities.

The goal of this manuscript was to understand to which extent was this year’s regional variation in burned area in southwest Europe extreme, relative to the 2001-2021 mean, and also to test the associated anomalies in terms of fuel moisture content and pyrometeorology.

## 2. Materials and methods

### 2.1. Fire data

Fire data for the 2000-2022 period were collected from the European Forest Fire Information System (EFFIS dataset; San-Miguel-Ayanz et al., 2012). We used EFFIS data instead of governmental records collected by each country as it represents an updated and harmonized data source at subcontinental scale. We employed data from the EFFIS real-time burned area and GlobFire databases, based on the MODIS Collection 6 (C6) MCD64A1 burned area product (Giglio et al., 2018). We retrieved daily burned area data over the full period, aggregating it at weekly level using the regional divisions by Calheiros et al. (2020) for Portugal, López Santalla & López García (2019) for Spain and Resco de Dios et al. (2021) for France.

### 2.2. Fuel moisture content and fire weather

We explored several indicators relating fuel moisture content and meteorological danger conditions. We examined trends in live fuel moisture content (LFMC) using a recently developed remotely-sensed product based on MODIS imagery (Cunill Camprubí et al., 2022). We also investigated temporal patterns in vapor pressure deficit (VPD), one of the main drivers of compounded live and dead fuel moisture content (Resco de Dios et al., 2021), following previous protocols (Nolan et al., 2016). Regarding fire weather, we explored the dynamics of the Hot-Dry-Windy (HDW) index at 925 hPa as formulated by Srock et al. (2018).

### 2.3. Statistical methods

We performed the interannual comparison of the cumulative distribution of burned area between different regions of southwestern Europe. Weekly data on area burned were aggregated into cumulative records during each year. The annual cumulative distributions were synthesized using 95% confidence intervals allowing the identification of anomalous seasons. The same procedure was replicated with LFMC, VPD and HDW data, but with non-cumulative data.

We also investigated the links between LFMC, VPD and HDW, and burned area during the summer season (June to August) by fitting linear regression models. To do so we aggregated each index as its period average and summarized the total burned area at yearly level. Individual models were trained for each region, reporting the slope of the regression line, the R^2^ and the significant level of the model (Fisher’s F test).

## 3. Results

### 3.1. The season in numbers

Burned area reached abnormally high values in some areas (Fig. 1). In southwest France, burned area between April-August 2022 exceeded the 2001-2021 median for the same period by a factor of 52 (27,228 ha in 2022 relative to 519.5 ha, Fig. 1f). Burned area in northwest (Fig. 1c) and in central Spain (Fig. 1d) also extended beyond historical records and it exceeded the 95^th^ percentile in southeast Spain (Fig. 1e). Burned area in northwest Portugal (Fig. 1a), southeast Portugal (Fig. 1b) and southeast France (Fig. 1g) was higher than the 2001-2021 median by a factor of 4, 2 and 5, respectively.

**Figure 1.**
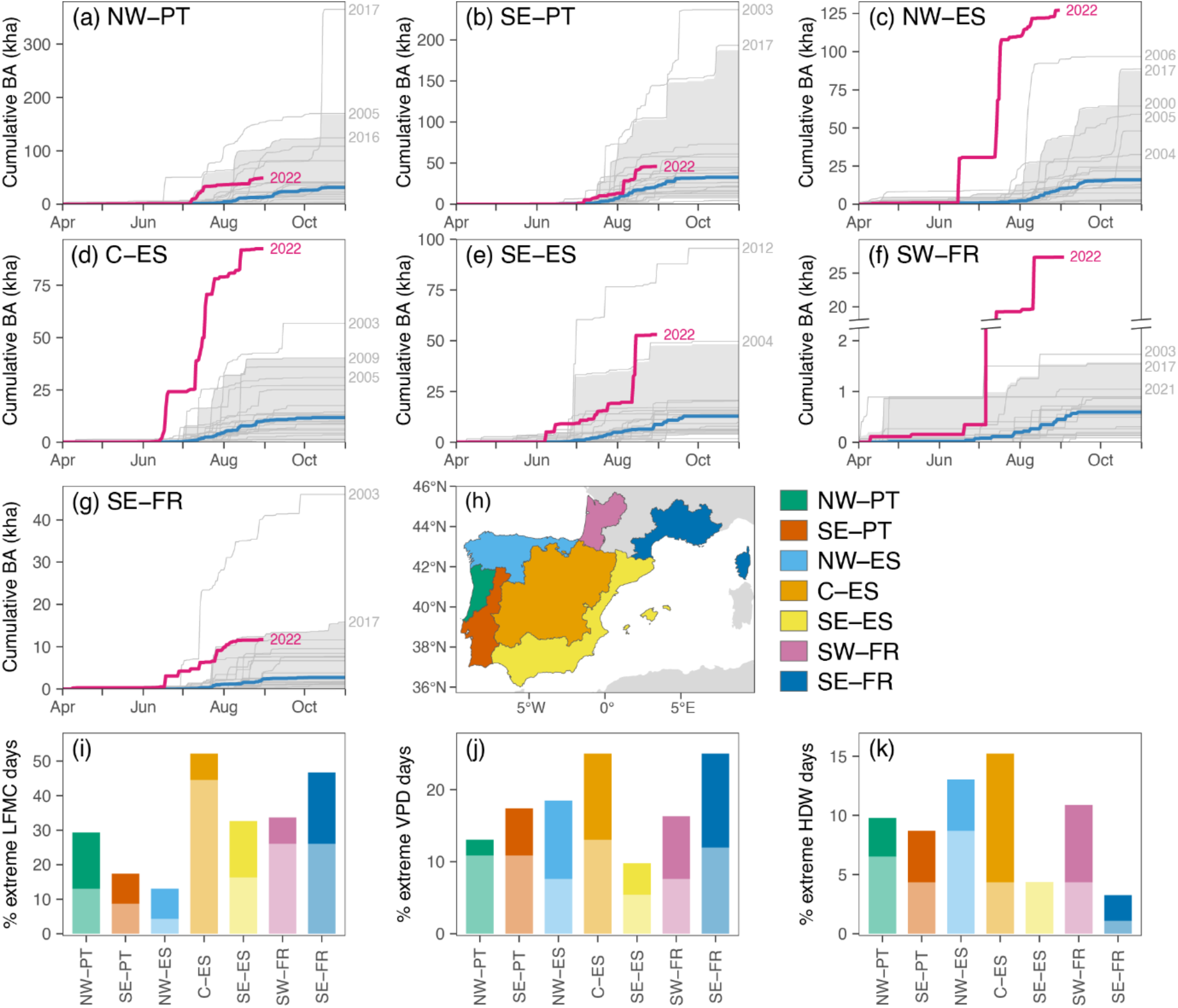
Cumulative burned area in SW Europe. The shaded area indicates the 95^th^ percentile in the historical records (2001-2021) and the blue and pink lines the historical median and current year, respectively. Panel i indicates the % of days between June-August when live fuel moisture content (LFMC) reached either record-breaking levels below the absolute minimum in the 2001-2021 registry (lower part of the bar) or below the driest 95^th^ quantile (upper part of the bar). Panels j and k indicate the % of days between June-August when vapor pressure deficit (VPD) and the hot-dry-weather index (HDW) reached either record-breaking levels above the absolute maximum in the 2001-2021 registry (lower part of the bar) or above the highest 95^th^ quantile (highest part of the bar graph). Data sources for panels i-k are provided in Figs. A1-A3.

Another anomalous situation of the 2022’s fire season was its early start, especially in southwest and southeast France and in central and northwest Spain. In those regions, large fires (>500 ha) are usually infrequent until mid to late August, whereas this year large fires occurred almost a month in advance (depending on region, Fig. 1). The onset of the season in the Spanish regions situates usually in July, though this year fire activity started in early-to-mid June. In France, the season started in April.

### 3.2. On the relationship with fuel moisture anomalies

Fuel dryness is a critical driver of fire ignition and spread in forested landscapes. LFMC reached record-breaking values during the 2022 season (Figs. 1, A1), ranging towards the driest historical conditions (Figs. 1i, A1). The major departure from the 2001-2021 median values (the driest anomaly) occurred in southwest France and central Spain, where 27% and 45% of the days between June-August coincided with values below the historical minimum of LFMC (Figs. 1i, A1), respectively. In the same line, we observed that 2022’s VPD values were towards the upper end, indicating a more desiccating atmosphere than under average conditions (Figs. 1j, A2). This season also showed record high VPD levels (relative to 2001-2021) during 10% of the fire season (June-August) across the entire area (Fig. 1j). The HDW index, despite depicting substantial day-to-day variability as expected in weather-related variables, denoted record high values for 5% of the fire season across southwest Europe (Figs. 1k, A3). Altogether, evidence suggests that dried than usual conditions have partly driven the extreme figures of the 2022’s season.

We observed significant effects in seasonal burned area of LFMC, VPD and HDW, though with regional differences (Table 1, Figs. A4-6). Northwest Portugal attained the strongest relationships with fuel moisture content (LFMC and VPD; R^2^=0.54 and 0.44, respectively), followed by central and northwest Spain, and southeast Portugal. The fire-HDW relationships were particularly strong in Portugal (northwest R^2^=0.61; southeast R^2^=0.37) and southeast France (R^2^=0.35). As expected, LFMC depicted a negative profile (lower moisture associated with higher burned area; Fig. A4) whereas VPD and HDW showed positive relationships (higher atmospheric drought or HDW associated with higher burned area; Figs. A5-A6). No significant relationship (p<0.05) was apparent for southeast Spain.

**Table 1.** Performance (R^2^) of the seasonal regression models of burned area during the summer season (June-August) against live fuel moisture content (LFMC), vapor pressure deficit (VPD) and the Hot-Dry-Windy index (HDW). Shading indicates the significance level of the relationships (dark grey, p<0.05; light grey, p<0.10).

|  | NW-PT | SE-PT | NW-ES | C-ES | SE-ES | SW-FR | SE-FR |
| --- | --- | --- | --- | --- | --- | --- | --- |
| LFMC | 0.54 | 0.25 | 0.24 | 0.28 | 0.01 | 0.10 | 0.31 |
| VPD | 0.44 | 0.11 | 0.32 | 0.40 | 0.05 | 0.19 | 0.39 |
| HDW | 0.61 | 0.37 | 0.13 | 0.08 | 0.04 | 0.00 | 0.35 |

## 4. Discussion

The evidence and data already available, although preliminary, clearly indicate that the 2022 wildfire season was extreme. Our findings revealed not only the extraordinary extent of wildfires, but also the early onset of the fire season associated with large fire events (Fig. 1). The implications and consequences of the shift toward extreme fire regimes are manifold and require careful consideration.

### 4.1. Fuel build-up, connectivity and aridity

The number of fires has declined in southwest Europe as a result of prevention campaigns and strong regulation over the last decades. The so-called “fire exclusion policy” has driven an overall decline in burned area in most Mediterranean countries (Rodrigues et al., 2020; Silva et al., 2019), creating a fire deficit that fosters fuel accumulation. Agricultural land abandonment and forest plantations have also contributed to landscape and fuel continuity, breaking the traditional protective land mosaic that once hindered fire spread.

Large fires require spatial connectivity of heavy fuel loads over landscape scales combined with fuel drying during protracted periods of water scarcity or heat waves. These conditions were met during the 2022’s summer months, with sustained high temperatures since May, leading to hazardous LFMC, VPD and HDW levels (Figs. 1i,j,k; Table 1; Figs. A1-3). The Copernicus Climate Service identified 2022 as an unusual year with exceptional heat wave events – in terms of frequency, intensity, and duration – striking the western Mediterranean Basin. However, this rare events fall inside the expected trend under climate warming projections and may even amplify over the next decades (C3S, 2017, 2022), potentially becoming average by 2035 (CCAG, 2022). Additionally, other factors related to the lack of fuel management (i.e. pyrosilviculture) in many forest stands is also likely to have contributed to the extreme burned area in the pine plantations that dominate the Landes, in southwest France, and other regions in central-northwest Spain and Portugal (Moreira et al., 2020).

### 4.2. An increased role of lightning-caused ignitions?

We still cannot examine full ignition causes as complete official records are not yet public or up-to-date. But anecdotal evidence and preliminary reports suggest that lightning was a major ignition source in northwest and southeast Spain (e.g., Sierra de la Culebra, ca. 33,000ha; Vall d’Ebo, ca. 12,000ha). Lightning is associated with more extreme fires as they occur under higher atmospheric instability (Fernandes et al., 2021). Over the Iberian Peninsula, lightning fires are known to be linked to dry thunderstorm episodes, particularly frequent in certain enclaves in the northwest of the Iberian Peninsula (Dijkstra et al., 2022). Summer thunderstorms are usually linked to thermal lows eventually developing after sustained anticyclonic conditions driving abnormally high temperatures (Fernandes et al., 2016; Rodrigues et al., 2019).

### 4.3. Implications and undesired impacts

Wildfire impacts are primarily social and economical in these fire-prone landscapes. That is, fire affects rural economies, and may favour further land abandonment as small-scale farming and forestry become less profitable. This may create a feedback loop, where fire enhances land abandonment, which then increases fuel connectivity and fuel loads and consequently further increases wildfire activity.

Earlier - and therefore potentially longer - seasons may have profound implications for forest and wildfire management as well. For instance, early onsets are likely to catch firefighting crews unprepared for a safe and efficient response, while preventive and management activities must also be scheduled sufficiently in advance.

Post-fire storms in recently burned areas may enhance erosion rates and soil losses could foster forest transformation into shrublands or grasslands, a transition that would bring increased fire spread potential (Scott & Burgan, 2005) and the loss of valuable ecosystem services (Morán-Ordóñez et al., 2021). Over large scales, fire-induced deforestation could lead to long-term land degradation, counteracting the increasing trend in forest area observed over the last decades (Karavani et al., 2018).

Fire suppression readily decreases burned area in the short-term, but the “fire suppression trap” implies that fuel accumulation resulting from oversuppression will increase burned area and the probability of extreme wildfires in the long-term. One could hypothesize that the year 2022 has been the turning point where, after decades of suppression-driven declining burned area, extreme wildfire seasons may increase from now on due to interactions between climate change and massive fuel accumulations. Although it is too early to test for this, it is clear that only landscape-scale fuel management can mitigate wildfire risk and break this reinforcing loop (Cochrane & Bowman, 2021; Moreira et al., 2020; Wunder et al., 2021).

## Acknowledgements

This work was supported by MICINN projects (RTI2018-094691-B-C31, PID2020-116556RA-I00); EU H2020 (grant agreements 101003890 and 101037419); and the Portuguese Foundation for Science and Technology (UIDB/04033/2020).

## Supplementary Information

**Fig. A1.**
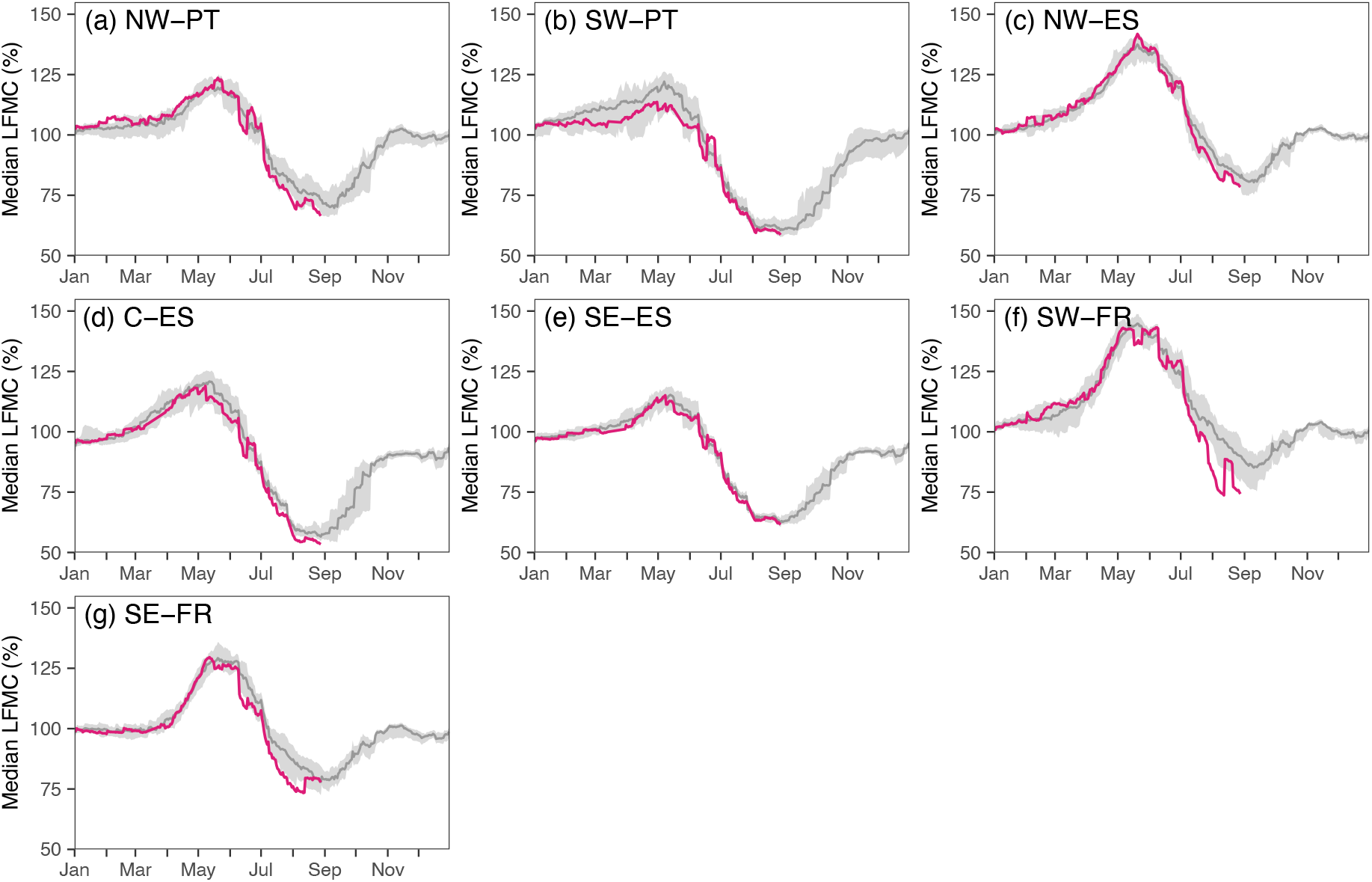
Temporal patterns in live fuel moisture content (LFMC) across the 7 regions of southwestern Europe from the data product developed by Cunill Camprubí et al. (2021). The red line indicates 2022 values while the grey line and shaded area denote the long-term (2001-2021) median and 95^th^ percentile.

**Fig. A2.**
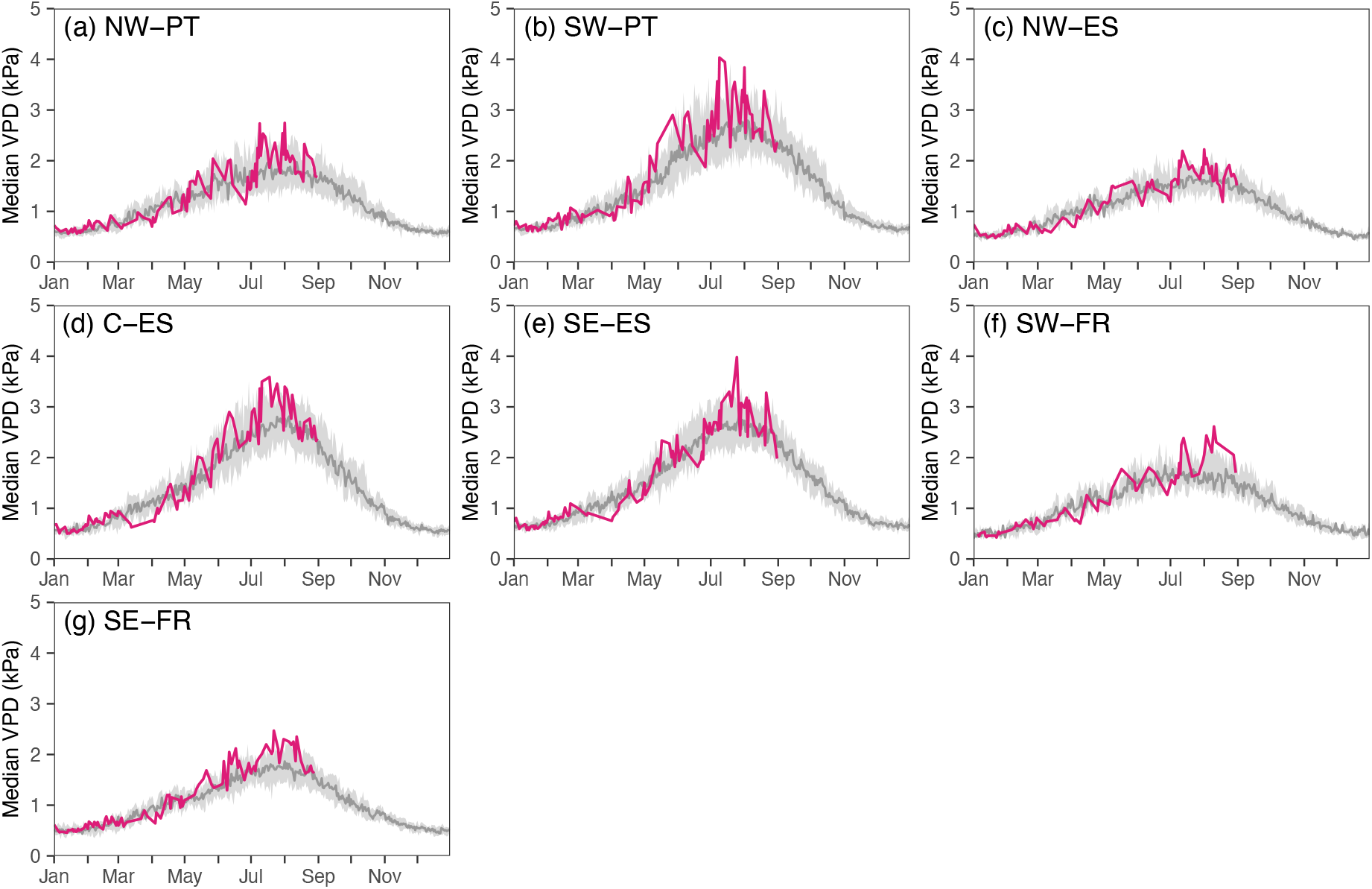
Temporal patterns in vapor pressure deficit (VPD) across the 7 regions of southwestern Europe following previous protocols (Nolan et al., 2016) and derived from MODIS LST MOD11A1 collection 6. The red line indicates 2022 values while the grey line and shaded area denote the long-term (2001-2021) median and 95^th^ percentile, respectively.

**Fig. A3.**
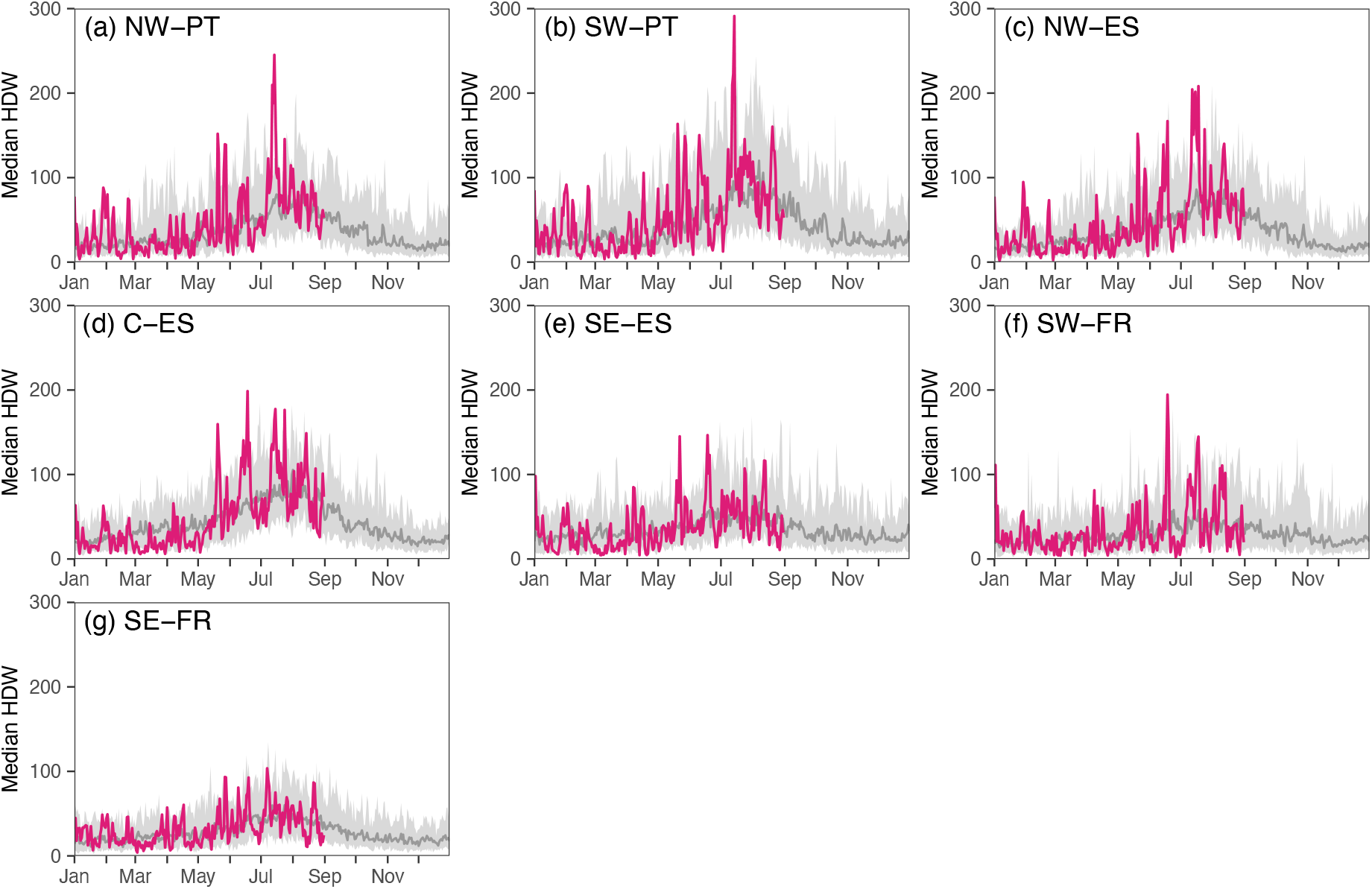
Temporal patterns in the hot-dry-windy index (HDW) at 925hPa across the 7 regions of southwestern Europe calculated following Srock et al., (2018) and using data from https://psl.noaa.gov/data/gridded/data.ncep.reanalysis2.html. The red line indicates 2022 values while the grey line and shaded area denote the long-term (2001-2021) median and 95^th^ percentile, respectively.

**Fig. A4.**
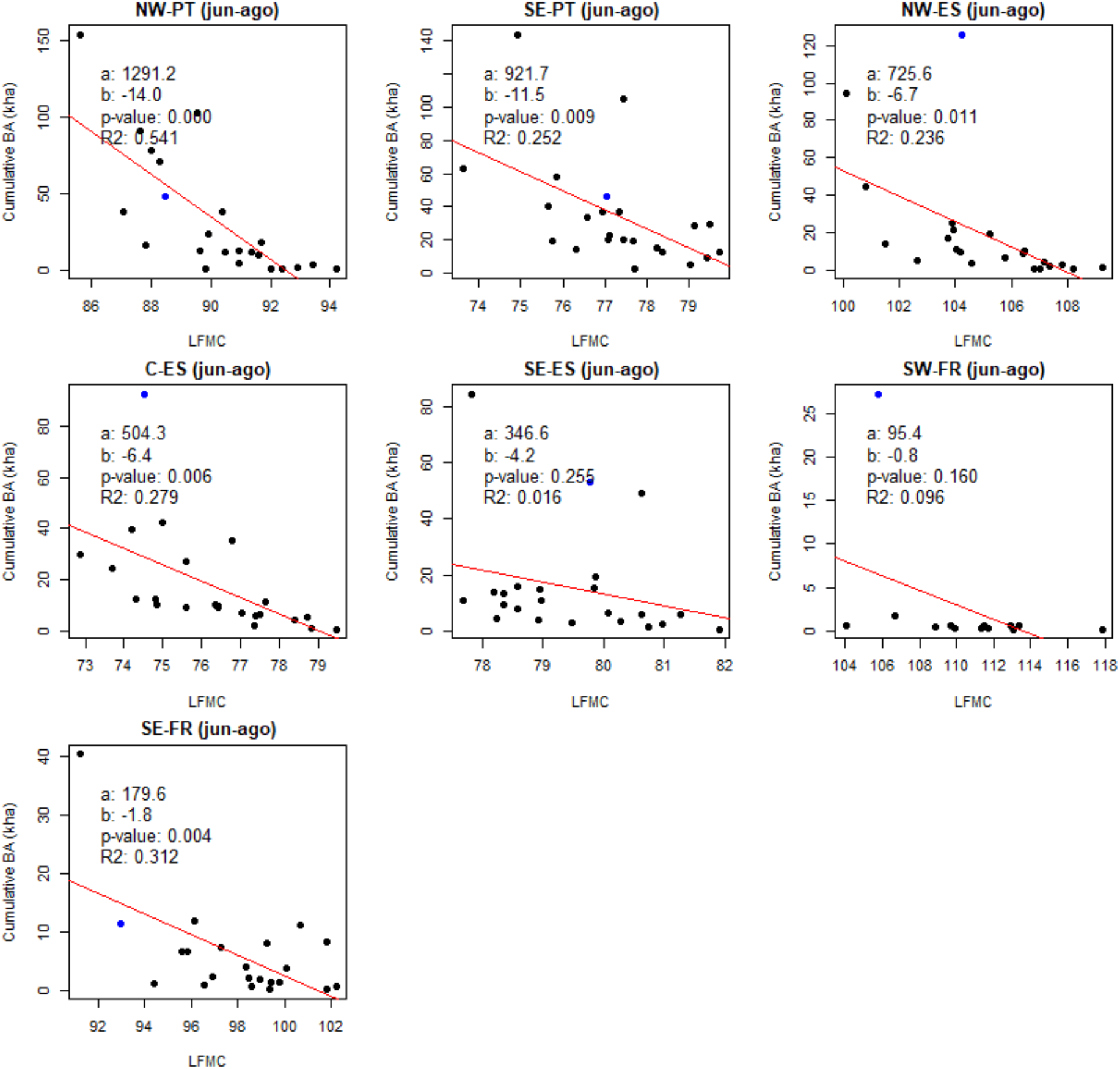
Season relationships between burned area and live fuel moisture content (LFMC) across the 7 regions of southwestern Europe from the model by Cunill Camprubí et al. (2021). a; intercept; b, slope of the regression line (in red); p-value, significance level; R2, Pearson’s coefficient of determination. The blue dot identifies the 2022 season.

**Fig. A5.**
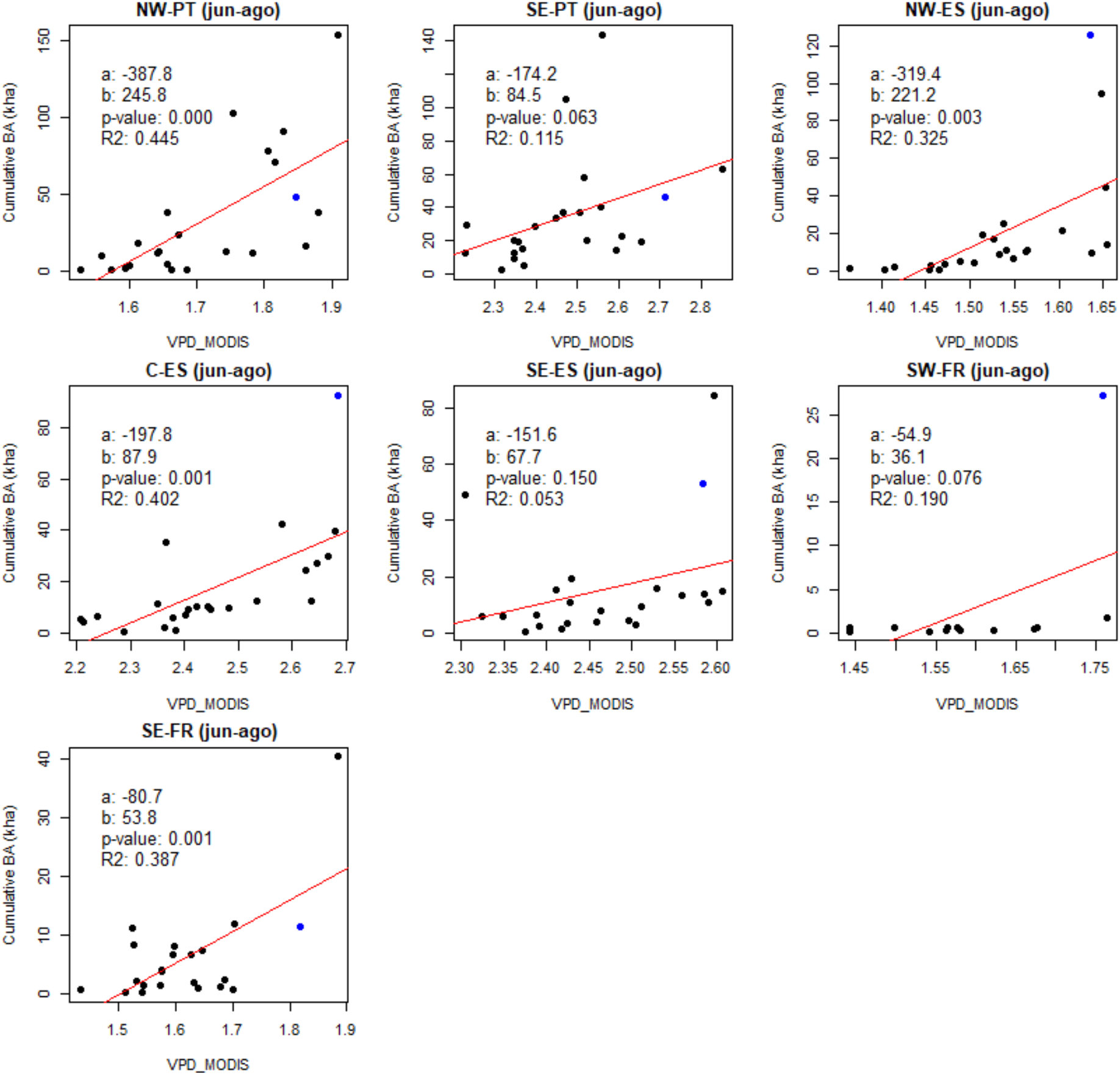
Season relationships between burned area and vapor pressure deficit (VPD) across the 7 regions of southwestern Europe following previous protocols (Nolan et al., 2016) and derived from MODIS LST MOD11A1 collection 6. a; intercept; b, slope of the regression line (in red); p-value, significance level; R2, Pearson’s coefficient of determination. The blue dot identifies the 2022 season.

**Fig. A6.**
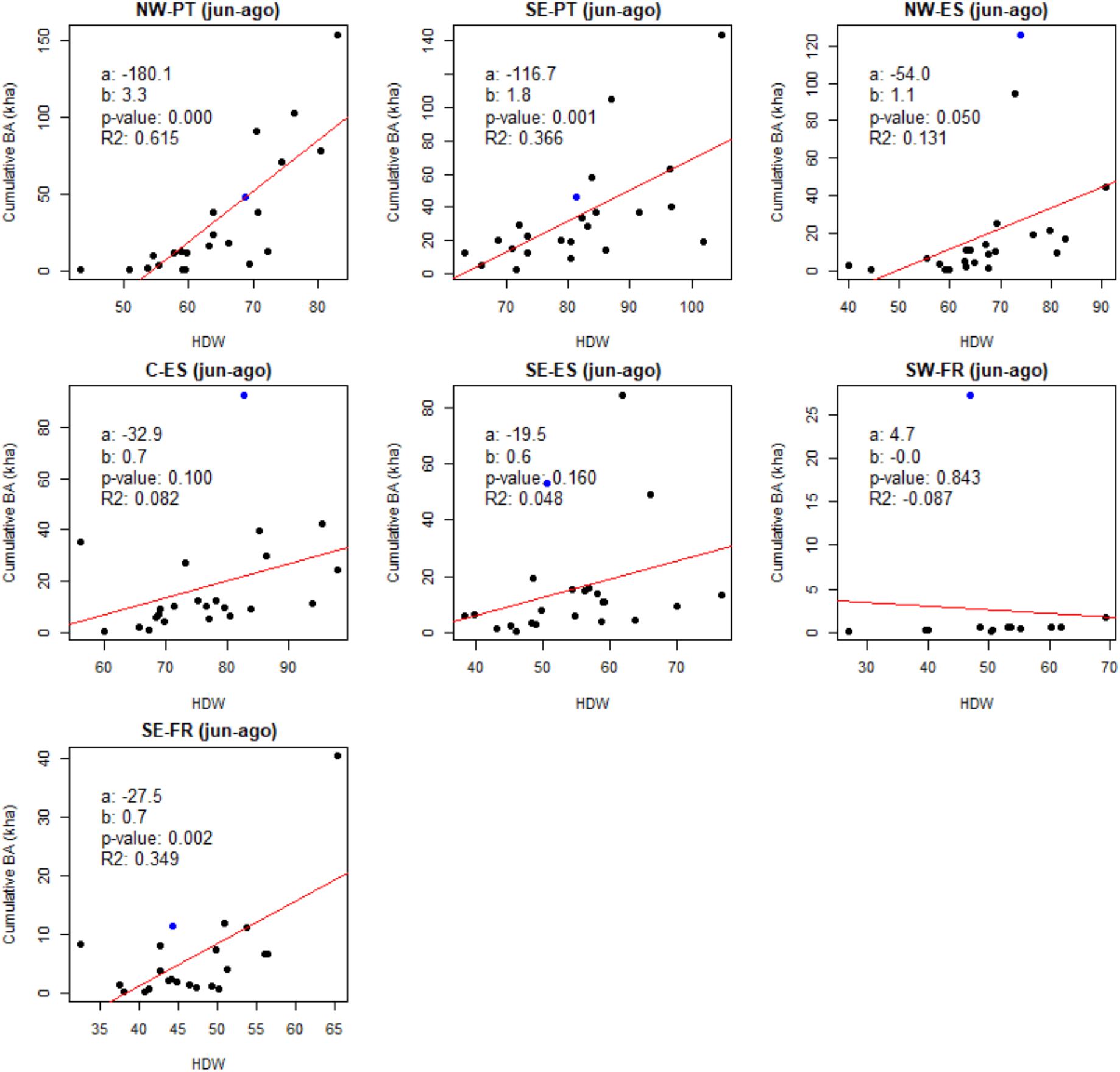
Season relationships between burned area and the Hot-Dry-Windy index (HDW) at 925hPa across the 7 regions of southwestern Europe calculated following Srock et al., (2018) and using data from https://psl.noaa.gov/data/gridded/data.ncep.reanalysis2.html. a; intercept; b, slope of the regression line (in red); p-value, significance level; R2, Pearson’s coefficient of determination. The blue dot identifies the season 2022.

